# Actin clearance promotes polarized dynein accumulation at the immunological synapse

**DOI:** 10.1101/505990

**Authors:** Elisa Sanchez, Xin Liu, Morgan Huse

**Author notes:** Equal contribution (Elisa Sanchez, Xin Liu). Department of Basic Medical Sciences, School of Medicine, Tsinghua University, Beijing 100084, China (Xin Liu).

## Abstract

Immunological synapse (IS) formation between a T cell and an antigen-presenting cell is accompanied by the reorientation of the T cell centrosome toward the interface. This polarization response is thought to enhance the specificity of T cell effector function by enabling the directional secretion of cytokines and cytotoxic factors toward the antigen-presenting cell. Centrosome reorientation is controlled by polarized signaling through diacylglycerol (DAG) and protein kinase C (PKC). This drives the recruitment of the motor protein dynein to the IS, where it pulls on microtubules to reorient the centrosome. Here, we used T cell receptor photoactivation and imaging methodology to investigate the mechanisms controlling dynein accumulation at the synapse. Our results revealed a remarkable spatiotemporal correlation between dynein recruitment to the synaptic membrane and the depletion of cortical filamentous actin (F-actin) from the same region, suggesting that the two events were causally related. Consistent with this hypothesis, we found that pharmacological disruption of F-actin dynamics in T cells impaired both dynein accumulation and centrosome reorientation. DAG and PKC signaling were necessary for synaptic F-actin clearance and dynein accumulation, while calcium signaling and microtubules were dispensable for both responses. Taken together, these data provide mechanistic insight into the polarization of cytoskeletal regulators and highlight the close coordination between microtubule and F-actin architecture at the IS.

## Introduction

Antigen recognition by the T cell receptor (TCR) induces the formation of a stereotyped interface between the T cell and the antigen-presenting cell known as the immunological synapse (IS) [1]. The IS maintains strong adhesion, regulates TCR signaling, and enables polarized intercellular communication. Critical to IS assembly is the reorientation of the T cell centrosome (also called the microtubule organizing center, or MTOC) to the center of the interface [2]. The centrosome polarizes to the IS within minutes of initial TCR stimulation, carrying with it the Golgi apparatus, endosomal compartment, and, in cytotoxic T cells, lytic granules containing perforin and granzyme. This promotes the directional secretion of cytokines and cytotoxic factors toward the APC, enhancing the specificity of these critical effector responses.

The centrosome is guided to the IS by cytoskeletal regulators that push and pull on the microtubule cytoskeleton [3-6]. Prior studies indicate that dynein, the predominant minus end-directed microtubule motor in eukaryotes, is particularly important for this process. Dynein accumulates at the IS shortly after TCR stimulation, and depletion or inhibition of the protein strongly impairs centrosome reorientation [4-7]. These results are consistent with a model in which dynein, after it becomes anchored at the synaptic membrane, pulls on microtubules to reposition the centrosome. How TCR signaling events modulate the localization of this molecular motor remains an area of active study. It is known that recruitment of dynein, and ultimately the centrosome, depends on the polarized accumulation of the lipid second messenger diacylglycerol (DAG) [7], which is generated via the hydrolysis of phosphatidylinostitol-4,5-bisphosphate (PIP_2_) by phospholipase C-γ (PLC-γ) downstream of the TCR. DAG forms a gradient centered at the IS, which drives cytoskeletal polarization, at least in part, by recruiting and activating members of the novel protein kinase C (nPKC) subfamily [8]. Precisely how DAG and nPKC signaling promotes recruitment of dynein, however, is poorly understood.

During IS assembly, cortical F-actin reorganizes into an annular configuration characterized by intense actin polymerization at the periphery of the contact and clearance from the center [1]. As with centrosome reorientation, formation of the synaptic F-actin ring depends strongly on phosphoinositide signaling. PIP_2_ depletion by PLC-γ promotes F-actin clearance at the center of the IS, while PIP_2_ phosphorylation by phosphoinositide 3-kinase drives F-actin growth in the periphery [9,10]. Furthermore, centrosome reorientation and F-actin ring formation are tightly coupled in time, with the centrosome moving to the IS just as F-actin clears from the central synaptic membrane [11]. These molecular and temporal relationships imply a close, and perhaps causal, relationship between F-actin remodeling and centrosome recruitment. Indeed, perturbations that disrupt F-actin ring formation also impair the centrosome polarization response [9,12]. Whether synaptic F-actin dynamics influence dynein localization, however, has not been explored.

In the present study, we applied a TCR photoactivation and imaging approach to investigate the mechanism of dynein localization to the IS. We found that F-actin depletion from the synaptic membrane was closely correlated with dynein accumulation in space and time, and that pharmacological inhibition of F-actin dynamics prevented both dynein recruitment and centrosome reorientation. These polarization responses were dependent on both DAG and PKC signaling, but did not require dynein activity or elevated cytoplasmic calcium (Ca^2+^). Taken together, these data establish the molecular and cellular framework for dynein polarization in T cells and provide insight into the assembly of immune cell-cell interactions.

## Results

### Localized TCR stimulation induces the recruitment of dynein 1 complexes

To study the recruitment dynamics of dynein in the context of polarized T cell activation, we employed an imaging-based method that enables spatiotemporally controlled stimulation of the TCR [13,14]. In this system, CD4^+^ T cells expressing the 5C.C7 TCR are added to glass surfaces coated with a photoactivatable form of their cognate ligand, the moth cytochrome c_88-_103 (MCC) peptide bound to the class II MHC protein I-E^k^ (MCC-I-E^k^). This photoactivatable peptide-MHC complex (called NPE-MCC-I-E^k^) is nonstimulatory to the 5C.C7 TCR until it is irradiated with ultraviolet (UV) light. After the T cells attach to the surface, focused UV irradiation is used to generate a micron-sized region of cognate peptide-MHC beneath an individual T cell, inducing localized TCR signaling responses and cytoskeletal remodeling events that can be monitored by epifluorescence or total internal reflection fluorescence (TIRF) microscopy. Using this system, we have found that TCR stimulation drives DAG gradient formation and the ordered recruitment of nPKC isoforms within 90 seconds [8]. This is followed by the localized accumulation of dynein ∼10 seconds later and the reorientation of the centrosome ∼5 seconds after that [7].

The precise subunit composition of cytoplasmic dynein varies depending on the cell type and intracellular compartment in question [15]. Using RT-PCR, we found that murine T cells express the full complement of cytoplasmic dynein 1 subunits with the exception of Dync1i1 (intermediate chain 1), which is found almost exclusively in the nervous system (Fig. 1A). To address the possibility that selective light intermediate and light chains are employed for cytoskeletal remodeling during IS formation, we expressed GFP-labeled forms of each subunit together with TagRFP-T-tubulin, a marker for the centrosome, in 5C.C7 effector T cells. Photoactivation studies revealed that both light intermediate chains of dynein 1 (Dync1li1 and Dync1li2), the roadblock light chains (Dylrb1 and Dylrb2), and the TcTex light chains (Dylt1b and Dylt3) accumulated in the region of TCR stimulation just prior to reorientation of the centrosome (white arrowheads in Fig. 1B). By contrast, the LC8 light chains were not recruited to the irradiated region. Instead, they appeared to associate with the centrosome itself and to move with it, consistent with previous work [16]. These results suggest that the pool of dynein 1 responsible for centrosome reorientation to the IS does not contain LC8 light chains but is otherwise unrestricted in its composition.

**Figure 1.**
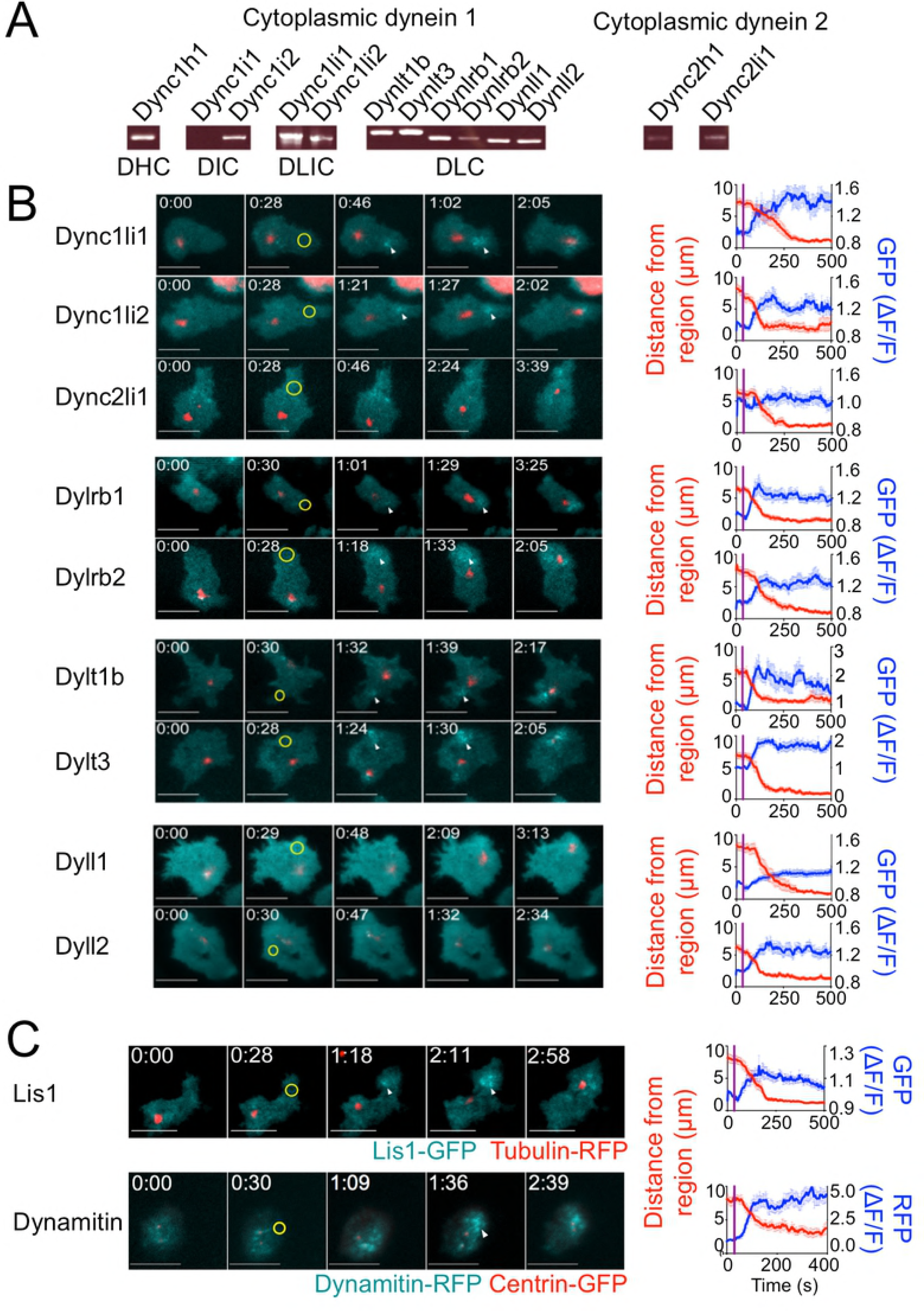
Cytoplasmic dynein 1 is recruited to the site of TCR stimulation. (A) RT-PCR analysis of the indicated dynein subunits, using RNA derived from 5C.C7 T cell blasts. (B-C) TCR photoactivation experiments were performed using 5C.C7 T cell blasts expressing fluorescently labeled tubulin or centrin (to visualize the centrosome) together with fluorescent probes for the indicated dynactin subunits (B) or dynein cofactors (C). Left, representative time-lapse montages showing TIRF images of the indicated dynein subunit or dynein cofactor and epifluorescence images of the centrosome. In each experiment, the moment and location of UV irradiation is indicated by a yellow circle. White arrowheads denote dynein or dynein cofactor accumulation in the plasma membrane prior to centrosome recruitment. Time in M:SS is indicated in the top left corner of each image. Right, graphs documenting the average dynein or dynein cofactor accumulation response (blue) and centrosome reorientation (red) in each data set. Error bars denote standard error of the mean (SEM). The moment of UV irradiation is indicated in each graph by a vertical purple line. Scale bars = 10 µm. In B, N ≥ 8 cells for each condition. In C, N ≥ 10 cells for each condition.

Although mature T cells do not possess primary cilia, they do express dynein 2 (Fig. 1A). However, we did not observe recruitment of the dynein 2 light intermediate chain (Dync2li1) to the IS upon photoactivation (Fig. 1B), suggesting that this motor complex is not involved in centrosome polarization in T cells. We also characterized the localization of two dynein cofactors, dynactin and Lis 1. Dynactin, which binds to dynein via its largest subunit p150^Glued^, regulates dynein’s motor processivity and its recruitment to microtubule plus ends [17]. Lis 1, for its part, directly binds to the motor domain of dynein and regulates its ability to move heavy cargoes [18]. To assess whether these cofactors are associated with synaptic dynein 1 during centrosome reorientation, we tagged Lis1 and the dynamitin subunit of dynactin with GFP and imaged them together with TagRFP-T-tagged tubulin in photoactivation experiments. Both Lis1 and dynactin were recruited to the irradiated region prior to centrosome reorientation (Fig. 1C), similar to the behavior we observed for dynein 1. Taken together, these results establish the relevant composition of dynein 1 complexes at the IS during TCR-induced polarization.

### TCR-induced F-actin clearance is required for dynein accumulation and centrosome reorientation

Having characterized the composition and dynamics of the synaptic dynein complex, we next investigated potential mechanisms for dynein recruitment. Previous studies have documented a correlation between F-actin clearance at the IS and centrosome polarization [5,11,19]. To investigate whether F-actin dynamics could modulate dynein accumulation, we performed TCR photoactivation experiments using 5C.C7 T cells expressing the F-actin probe Lifeact-mRuby2 together with Dync1li1-GFP (to monitor dynein). Accumulation of dynein in the irradiated region was markedly associated with the depletion of cortical F-actin (Figure 2A). This anti-correlation was particularly striking in cells that exhibited multiple cycles of Lifeact-mRuby2 depletion and recovery; in each cycle, the Dync1li1-GFP probe displayed inverse behavior (magenta arrows in Fig. 2A).

**Figure 2.**
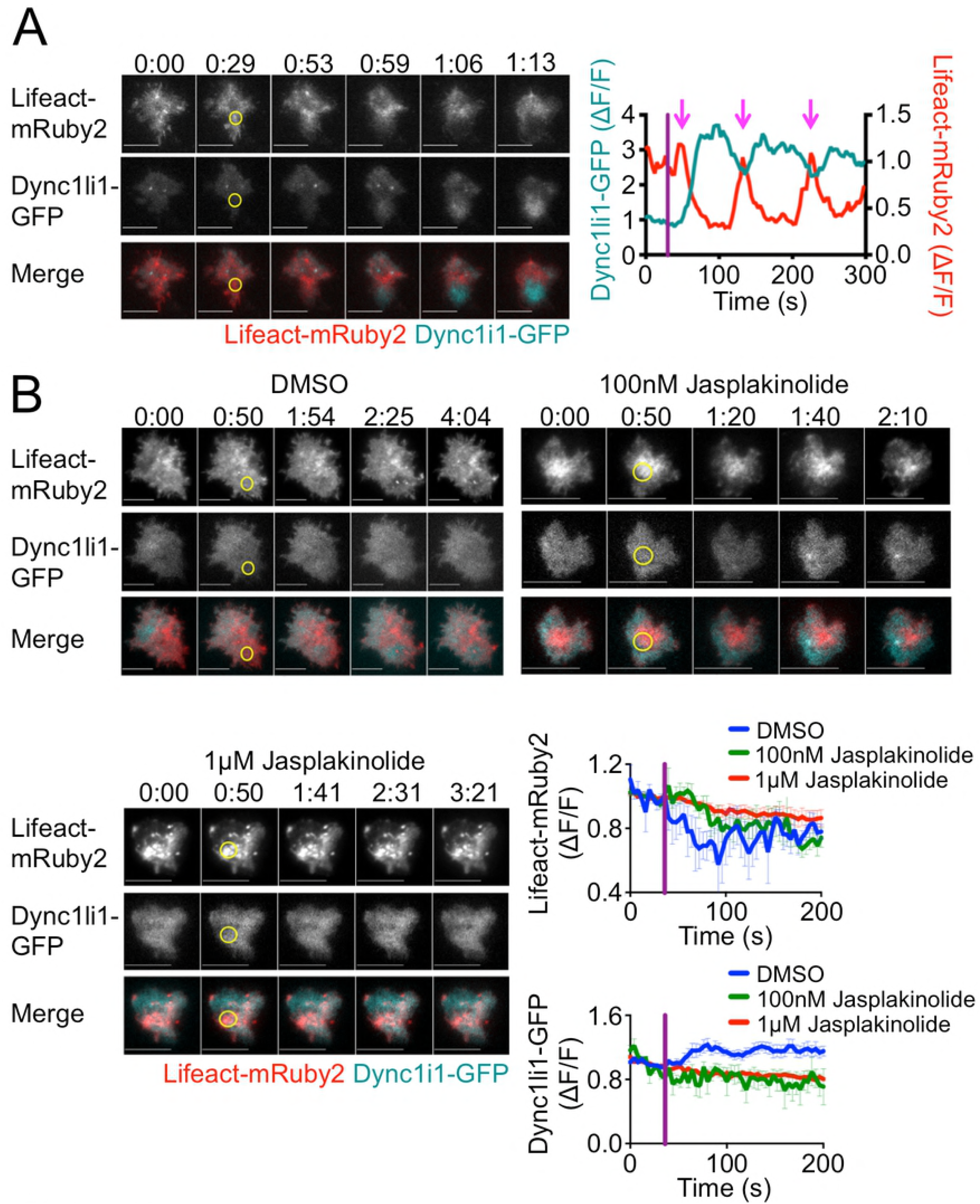
TCR-induced dynein accumulation requires F-actin depletion. TCR photoactivation experiments were performed using 5C.C7 T cell blasts expressing Dync1li1-GFP together with Lifeact-mRuby2 in the presence or absence of jasplakinolide. (A) A representative time-lapse montage is shown on the left, with the corresponding F-actin (red) and dynein (cyan) responses graphed to the right. Magenta arrows mark the cycles of F-actin depletion and dynein accumulation exhibited by this cell. (B) Top and bottom left, representative time-lapse montages showing TIRF images of F-actin and dynein in the presence of the indicated amounts of jasplakinolide or vehicle control (DMSO). Bottom right, graphs showing mean actin depletion (top) and dynein accumulation (bottom) in the presence of the indicated amounts of jasplakinolide or vehicle control (DMSO). In time-lapse montages, the moment and location of UV irradiation is indicated by a yellow circle. Time in M:SS is indicated above each montage. Scale bars = 10 µm. In graphs, error bars denote SEM, and vertical purple lines indicate the moment of UV irradiation. N ≥ 10 cells for each condition.

These results suggested a link between cortical F-actin and dynein recruitment at the IS. To explore this relationship, we carried out photoactivation experiments in the presence of jasplakinolide, a small molecule that promotes the polymerization and stabilization of F-actin. Jasplakinolide profoundly inhibited both Lifeact-mRuby2 clearance as well as Dync1li1-GFP accumulation at the irradiated region, consistent with the idea that F-actin depletion is a requisite step for the recruitment of dynein (Fig. 2B). To probe the downstream consequences of this disruption in dynein, we examined centrosome reorientation in jasplakinolide-treated cells. Centrosome recruitment to the irradiated region was inhibited by jasplakinolide in a dose dependent manner (Fig 3A), indicating that F-actin clearance is necessary for T cell polarization. Notably, the effect of jasplakinolide on dynein accumulation was markedly stronger than its effect on centrosome reorientation. This could be due to compensation by myosin II, which also contributes to centrosome movement and may be less sensitive to F-actin stabilization [5].

**Figure 3.**
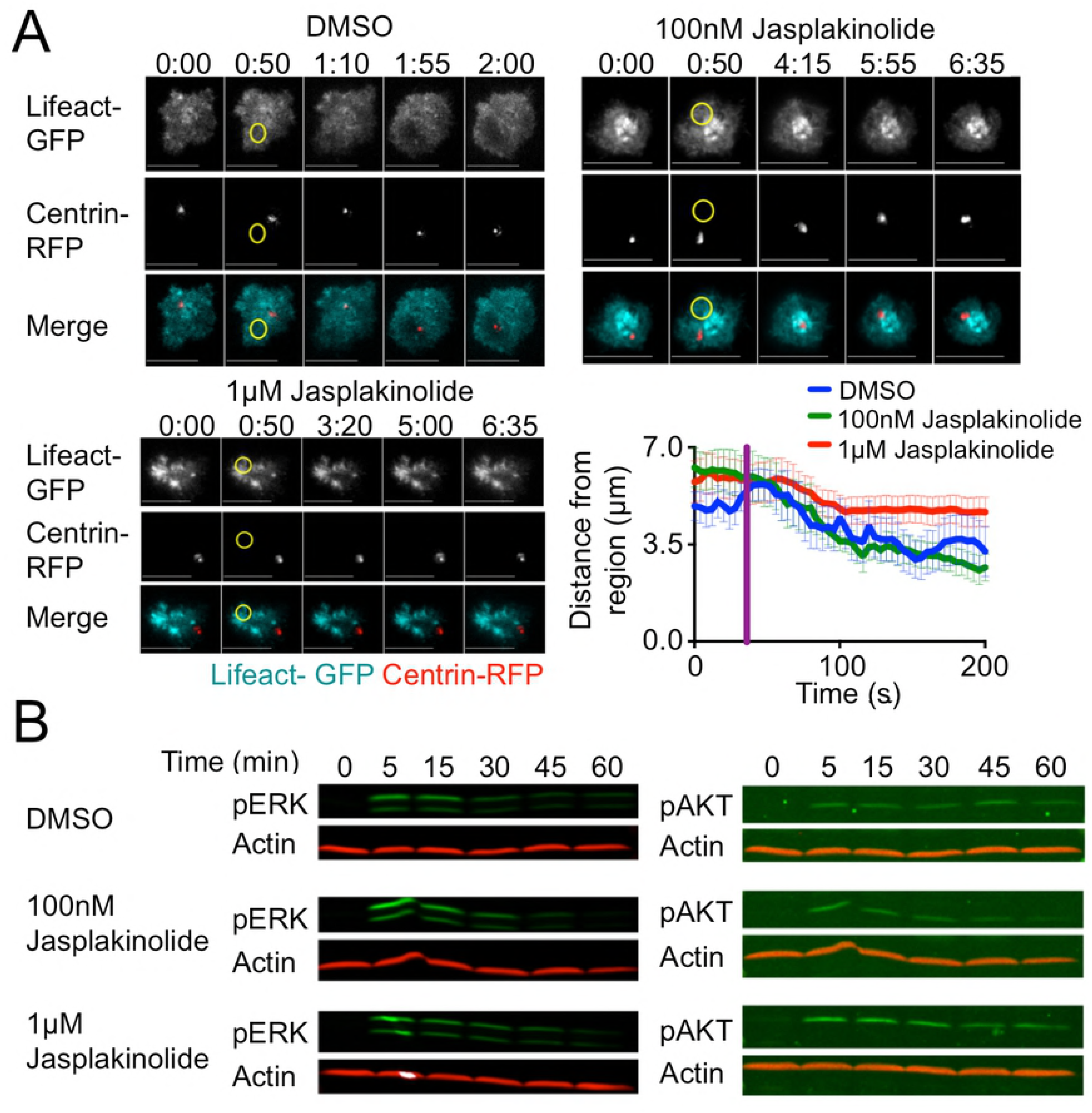
F-actin clearance promotes centrosome reorientation. (A) TCR photoactivation experiments were performed using 5C.C7 T cell blasts expressing Lifeact-GFP together with Centrin-TagRFP-T (Centrin-RFP) in the presence of the indicated amounts of jasplakinolide or vehicle control (DMSO). Top and bottom left, representative time-lapse montages showing TIRF images of F-actin and epifluorescence images of the centrosome. The moment and location of UV irradiation is indicated by a yellow circle. Time in M:SS is indicated above each montage. Scale bars = 10 µm. Bottom right, centrosome reorientation was quantified as the mean distance between the centrosome and the center of the irradiated region over time. Error bars denote SEM, and the vertical purple line indicates the moment of UV irradiation. N ≥ 8 cells for each condition. (B) 5C.C7 T cell blasts were stimulated using beads coated with MCC-I-E^k^ and ICAM-1 in the presence of the indicated amounts of jasplakinolide or vehicle control (DMSO). pERK1/2 and pAKT responses were assessed at the indicated times by immunoblot.

Although these results were consistent with a specific role for F-actin clearance in the recruitment of dynein, we also considered the possibility that jasplakinolide altered dynein accumulation secondarily by globally inhibiting TCR signaling. 5C.C7 T cells were preincubated with jasplakinolide or vehicle and then mixed with polystyrene beads coated with MCC-I-E^k^ and the adhesion protein ICAM-1, a ligand for the α_L_β_2_ integrin LFA-1. TCR-induced activation of the MAP kinase and phosphoinositide 3-kinase pathways was then assessed by immunoblot for phospho-Erk1/2 and phospho-AKT, respectively. Jasplakinolide did not alter these signaling outputs (Fig. 3B), implying that its effects on dynein recruitment and centrosome reorientation reflected a specific role for F-actin dynamics in both of these processes.

### Localized PKC activity is required for F-actin clearance and dynein recruitment

Previous work has established the importance of polarized DAG signaling for centrosome reorientation and dynein recruitment [7]. Accordingly, we investigated whether DAG accumulation is involved in F-actin depletion at the IS. To disrupt the effects of polarized DAG, we treated 5C.C7 T cells expressing Lifeact-mRuby2 and Dync1li1-GFP with phorbol 12,13-dibutyrate (PDBU), a phorbol ester that induces unpolarized DAG signaling. In photoactivation experiments, PDBU treatment disrupted both F-actin clearance and dynein accumulation at the irradiated region, indicating that polarized DAG is indeed necessary for both processes (Fig. 4A).

**Figure 4.**
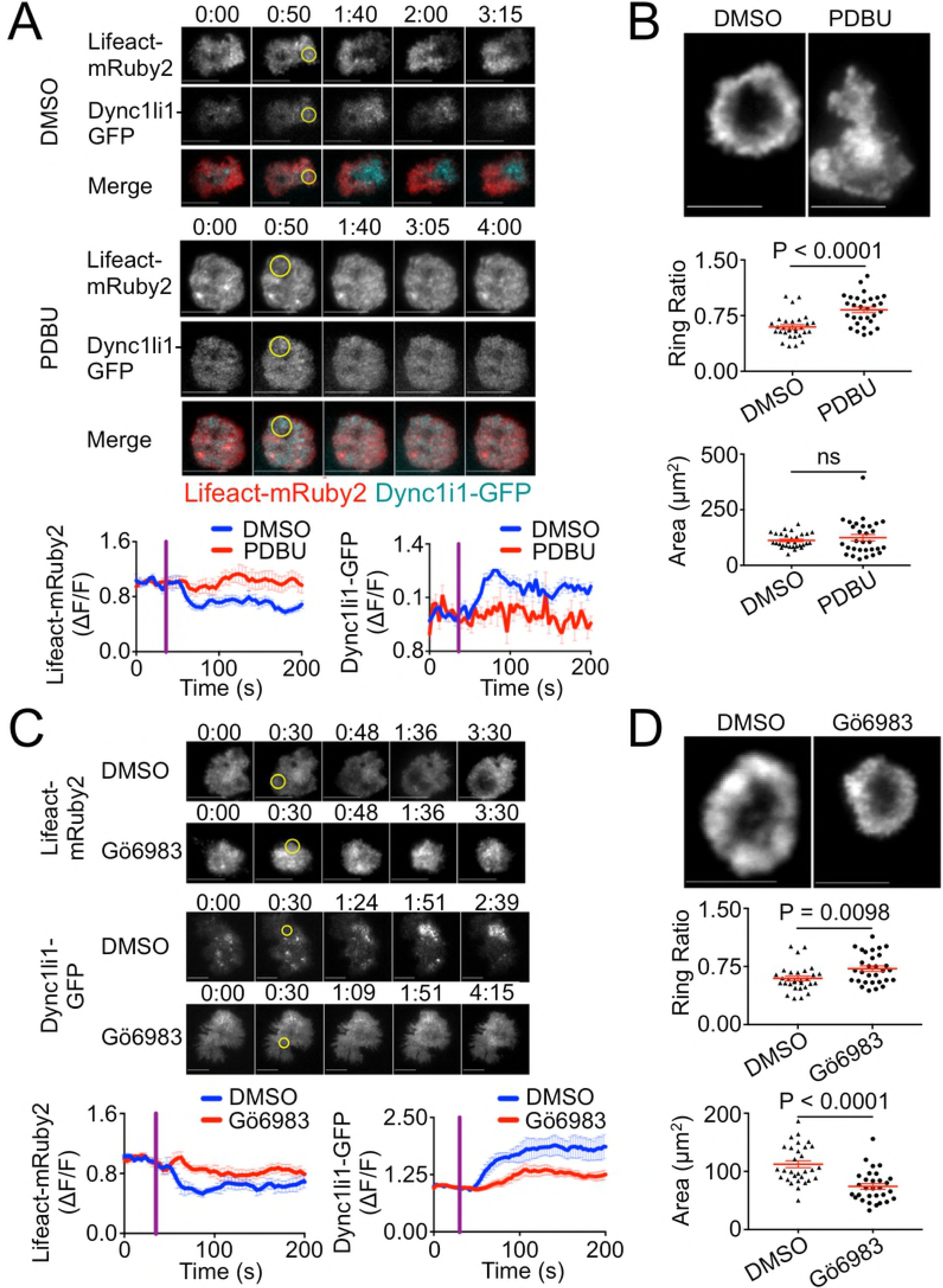
Polarized DAG and PKC signaling control synaptic F-actin clearance. (A) TCR photoactivation experiments were performed using 5C.C7 T cell blasts expressing Lifeact-mRuby2 together with Dync1li1-GFP in the presence of 1 µM PDBU or vehicle control (DMSO). Above, representative time-lapse montages showing TIRF images of F-actin and dynein. The moment and location of UV irradiation is indicated by a yellow circle. Time in M:SS is indicated above each montage. Scale bars = 10 µm. Below, quantification of mean F-actin clearance (left) and dynein accumulation (right) over time. Error bars denote SEM, and vertical purple lines indicate the moment of UV irradiation. N ≥ 13 cells for each condition. (B) 5C.C7 T cell blasts were stimulated on supported lipid bilayers containing MCC-I-E^k^ and ICAM-1 in the presence of 1 µM PDBU or vehicle (DMSO), fixed, and stained with phalloidin to visualize F-actin. Above, TIRF images of representative synapses. Below, quantification of ring ratio (top) and IS area (bottom). Mean values and error bars (SEM) are shown in red. N = 30 cells for each condition. (C) TCR photoactivation experiments were performed using 5C.C7 T cell blasts expressing Lifeact-mRuby2 together with Dync1li1-GFP in the presence of 50 nM Gö6983 or vehicle control (DMSO). Above, representative time-lapse montages showing TIRF images of F-actin and dynein. Time in M:SS is indicated above each montage. Scale bars = 10 µm. Below, quantification of mean F-actin clearance (left) and dynein accumulation (right) over time. Error bars denote SEM, and vertical purple lines indicate the moment of UV irradiation. N ≥ 6 cells for each condition. (D) 5C.C7 T cell blasts were stimulated on supported lipid bilayers containing MCC-I-E^k^ and ICAM-1 in the presence of 50 nM Gö6983 or vehicle (DMSO), fixed, and stained with phalloidin to visualize F-actin. Above, TIRF images of representative synapses. Below, quantification of ring ratio (top) and IS area (bottom). Mean values and error bars (SEM) are shown in red. N = 30 cells for each condition. P-values calculated from two-tailed unpaired Student’s T-test.

To further assess DAG-mediated F-actin clearance at the IS, we imaged PDBU-treated 5C.C7 T cells on supported lipid bilayers coated with stimulatory MCC-I-E^K^ and ICAM-1. T cells typically form radially symmetric synapses on these bilayers containing a peripheral F-actin ring and a central domain that is depleted of F-actin (Fig. 4B). In the presence of PDBU, however, T cells displayed asymmetric patterns without obvious F-actin clearance in the center (Fig. 4B). To quantify this organizational defect, we used the “ring ratio” parameter, which compares the fluorescence intensity at the periphery of the IS with that of the center [10]. A ring ratio less than one is indicative of an annular fluorescence pattern, whereas a ring ratio of one denotes a uniform distribution. PDBU treatment led to a substantial increase in ring ratio (Fig. 4B), consistent with profound disruption of F-actin clearance. Despite these dramatic effects on F-actin configuration, PDBU did not significantly alter IS size (Fig. 4B). Hence, the polarized delivery of DAG signals is crucial for IS organization, but it appears to be unnecessary for the formation of the contact itself.

DAG drives centrosome reorientation and dynein recruitment to the IS at least in part through nPKCs [8]. To explore the role of PKCs in F-actin clearance, we performed photoactivation experiments using 5C.C7 T cells pretreated with the PKC inhibitor Gö6983. F-actin clearance and dynein accumulation at the irradiated region were both attenuated by Gö6983 (Fig. 4C), consistent with a role for PKC activity in both processes. Gö6983 also inhibited F-actin ring formation on stimulatory lipid bilayers (Fig. 4D), although its effects were less dramatic than those of PDBU. Interestingly, Gö6983 treated T cells formed significantly smaller synapses than vehicle treated controls (Fig. 4D), demonstrating that insufficient PKC activity can stunt IS growth. Collectively, these results indicate that polarized DAG and PKC signaling control the growth and organization of F-actin structures at the IS.

### Calcium signaling is dispensable for TCR-induced F-actin clearance

Previous work has suggested that calcium (Ca^2+^) flux is required for F-actin clearance at the IS [20]. To examine whether Ca^2+^ signaling controls F-actin depletion and dynein recruitment during centrosome polarization, we performed TCR photoactivation experiments using T cells expressing Lifeact-mRuby2 and Dync1li1-GFP in medium containing the Ca^2+^ chelator EGTA. Surprisingly, F-actin clearance from the irradiated region was unaffected by the absence of Ca^2+^ (Fig. 5A). Dynein accumulation was also normal, reaffirming the close relationship between these two processes. To further assess the importance of Ca^2+^ signaling for synaptic F-actin clearance, we incubated T cells on stimulatory lipid bilayers under Ca^2+^-free and control conditions and imaged the resulting synapses. In agreement with the photoactivation experiments, we observed robust F-actin clearance from the center of the IS both in the presence and the absence of extracellular Ca^2+^ (Fig. 5B). Furthermore, Ca^2+^ depletion did not significantly alter IS size. Collectively, these results indicate that Ca^2+^ signaling does not strongly affect synaptic F-actin architecture and dynein recruitment.

**Figure 5.**
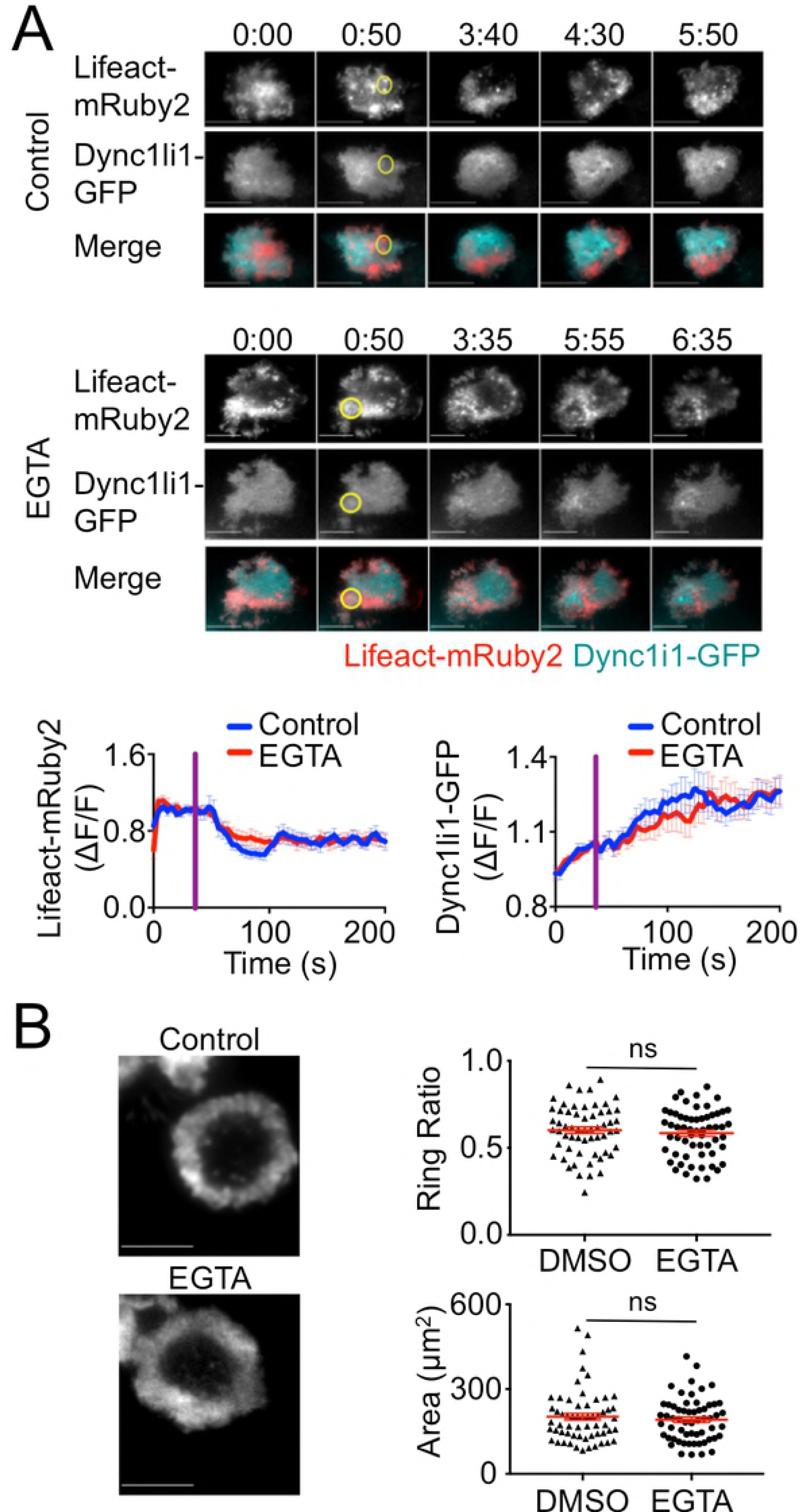
Calcium signaling is dispensable for F-actin clearance. (A) TCR photoactivation experiments were performed using 5C.C7 T cell blasts expressing Lifeact-mRuby2 together with Dync1li1-GFP in the presence or absence of 1 mM EGTA with 4 mM MgCl_2_ (to deplete extracellular Ca^2+^). Above, representative time-lapse montages showing TIRF images of F-actin and dynein. The moment and location of UV irradiation is indicated by a yellow circle. Time in M:SS is indicated above each montage. Scale bars = 10 µm. Below, quantification of mean F-actin clearance (left) and dynein accumulation (right) over time. Error bars denote SEM, and vertical purple lines indicate the moment of UV irradiation. N ≥ 14 cells for each condition. (B) 5C.C7 T cell blasts were stimulated on supported lipid bilayers containing MCC-I-E^k^ and ICAM-1 in the presence or absence of 1 mM EGTA with 4 mM MgCl_2_, fixed, and stained with phalloidin to visualize F-actin. Left, TIRF images of representative synapses. Right, quantification of ring ratio (top) and IS area (bottom). Mean values and error bars (SEM) are shown in red. N = 59 cells for each condition. P-values calculated from two-tailed unpaired Student’s T-test.

### Microtubules and dynein activity are dispensable for TCR-induced dynein recruitment

Microtubules are thought to deliver dynein to the plasma membrane in other cell types [21,22]. To examine whether a similar mechanism mediates dynein recruitment to the IS, we applied small molecule modulators of the microtubule cytoskeleton. 5C.C7 T cells expressing Lifeact-mRuby2 and Dync1li1-GFP were treated with either the microtubule depolymerizing agent nocodazole or the stabilizing agent taxol and then subjected to TCR photoactivation. Neither nocodazole nor taxol affected TCR-induced F-actin dynamics (Fig. 6A). In both cases, clearance of Lifeact-mRuby2 from the irradiated region was essentially indistinguishable from that of control cells. Nocodazole-treated and taxol-treated T cells also exhibited robust dynein recruitment to the irradiated region (Fig. 6A). Indeed, in the case of taxol, Dync1li1-GFP accumulation was consistently more intense than in cells treated with vehicle alone. Taken together, these results strongly suggest that microtubules are not involved in dynein delivery to the IS. Importantly, both nocodazole and taxol inhibited centrosome reorientation in photoactivation experiments (Fig. 6B). Hence, while microtubules are required for cytoskeletal polarization, they operate at a step downstream of motor protein recruitment.

**Figure 6.**
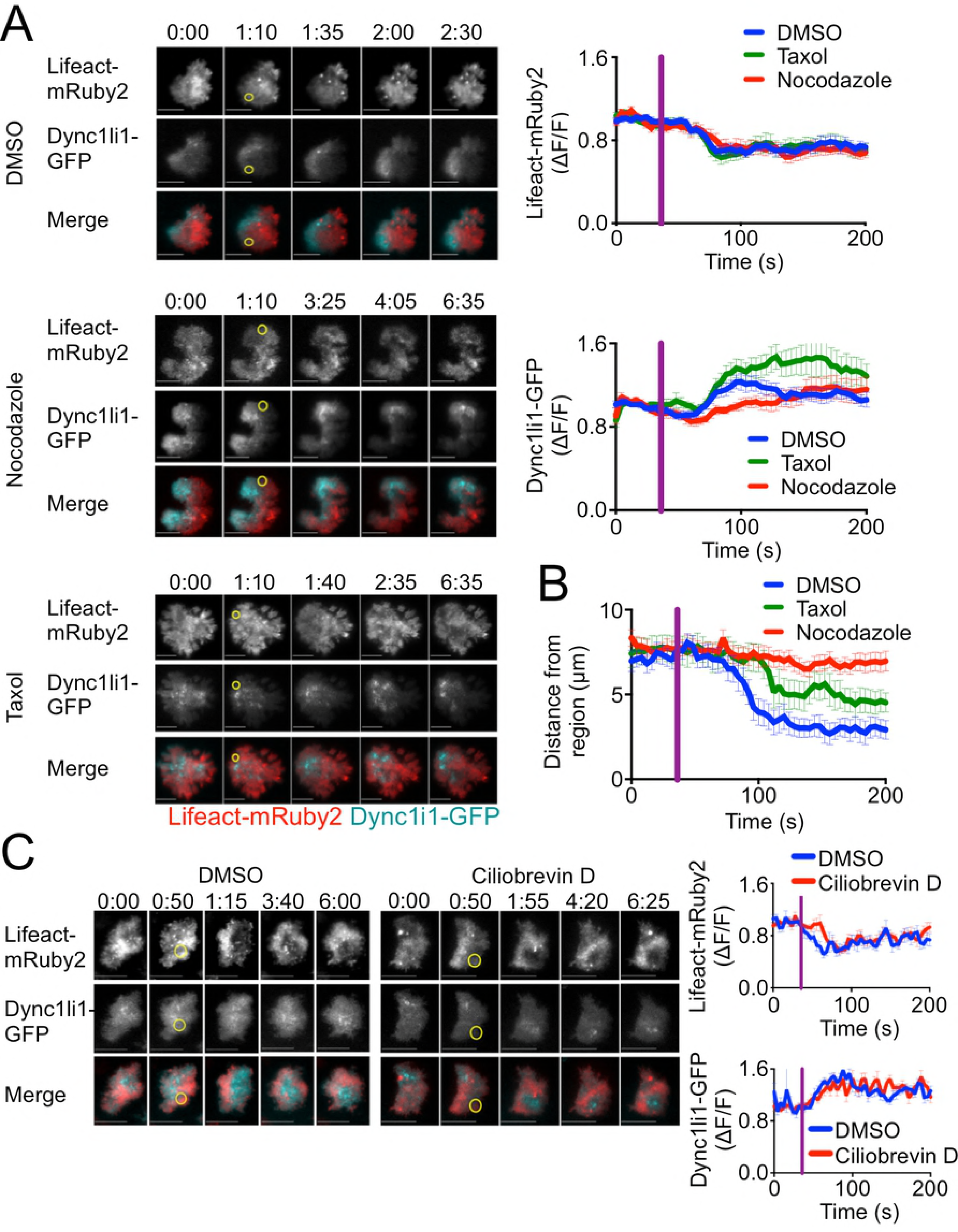
Microtubules and dynein activity are dispensable for F-actin depletion and dynein recruitment. (A) TCR photoactivation experiments were performed using 5C.C7 T cell blasts expressing Lifeact-mRuby2 together with Dync1li1-GFP in the presence of 30 µM nocodazole, 1 µM taxol, or vehicle control (DMSO). Left, representative time-lapse montages showing TIRF images of F-actin and dynein. Right, quantification of mean F-actin clearance (top) and dynein accumulation (bottom) over time. N ≥ 14 cells for each condition. (B) TCR photoactivation experiments were performed using 5C.C7 T cell blasts expressing Centrin-TagRFP-T in the presence of 30 µM nocodazole, 1 µM taxol, or vehicle control (DMSO). Centrosome reorientation was quantified by determining the mean distance between the centrosome and the center of the irradiated region over time. N ≥ 14 cells for each condition. (C) TCR photoactivation experiments were performed using 5C.C7 T cell blasts expressing Lifeact-mRuby2 together with Dync1li1-GFP in the presence of 50 µM ciliobrevin D or vehicle control (DMSO). Left, representative time-lapse montages showing TIRF images of F-actin and dynein. Right, quantification of mean F-actin clearance (top) and dynein accumulation (bottom) over time. N ≥ 9 cells for each condition. In all time-lapse montages, the moment and location of UV irradiation is indicated by a yellow circle. Time in M:SS is indicated above each montage. Scale bars = 10 µm. In all graphs, error bars denote SEM, and vertical purple lines indicate the moment of UV irradiation.

Finally, we examined whether the motor activity of dynein is necessary for its own recruitment. 5C.C7 T cells expressing Lifeact-mRuby2 and Dync1li1-GFP were subjected to TCR photoactivation in the presence of ciliobrevin D, a small molecule dynein inhibitor. TCR-induced F-actin clearance and dynein accumulation were unaffected by ciliobrevin D (Fig. 6C), indicating that motor activity is not required for dynein accumulation at the IS.

## Discussion

Proper IS function relies not only upon robust F-actin polymerization at the cell-cell interface but upon its targeted depletion, as well. Cortical F-actin clearance within the IS plays a well-established role in the directional secretion of cytotoxic molecules toward the target cell [23,24], and it has also been linked to the reorientation of the centrosome [11,19]. Here, we demonstrate the importance of F-actin depletion for the synaptic accumulation of dynein, a microtubule motor that is critical for both T cell polarity and intracellular trafficking. This work is consistent with the idea that F-actin and microtubule dynamics are closely coupled at the IS [25], which is fitting for a structure that mediates highly coordinated exo- and endocytosis.

The diversity of dynein isoforms, light chain subunits, and cofactors presumably facilitates functional specialization of the motor in different intracellular contexts. Our imaging experiments suggest that the synaptic pool of dynein involved in centrosome polarization contains the intermediate and light intermediate chains of cytoplasmic dynein1 along with the roadblock and Tctex light chains. The LC8 light chain, by contrast, did not accumulate in the region of TCR stimulation, although it did appear to decorate centrosome-associated compartments. These results imply a division of labor between the dynein light chains at the IS, with LC8 contributing to organelle accumulation at the centrosome but not centrosome reorientation per se. Notably, we did not observe synaptic recruitment of cytoplasmic dynein2, the isoform associated with intraflagellar transport at cilia [15]. Recent studies have highlighted the structural and molecular similarities between the IS and primary cilia [26,27]; dynein usage does not appear to be one of them. We also observed accumulation of dynamitin and Lis1 in the region of TCR stimulation, implying that both the dynactin and the Lis1/NDE1 complexes are involved in IS assembly. The precise role of dynactin in T cell polarity is somewhat controversial. Although overexpression of the dynamitin subunit was initially shown to impair centrosome reorientation [4], recent shRNA knockdown studies have suggested that it is Lis1/NDE1, and not dynactin, that operates in this context [28]. Although our data are consistent with roles for both complexes, they do not exclude the possibility that dynactin may associate with the synaptic dynein complex without contributing to its function.

Blocking TCR-induced Ca^2+^ signaling had no effect on F-actin clearance and centrosome reorientation in our hands. These results, although consistent with our previous work [7], contradict a recent report indicating that Ca^2+^ is required for synaptic F-actin depletion [20]. Whereas we employed Lifeact to label F-actin in our live imaging experiments, the previous study used F-tractin, which is thought to label a larger fraction of filaments [29]. Hence, it is formally possible that we failed to observe a Ca^2+^ sensitive pool of F-actin at the IS that could be detected by F-tractin but not by Lifeact. Incomplete Lifeact labeling, however, does not explain our phalloidin staining experiments, which also failed to support a role for Ca^2+^ signaling. Furthermore, even if a Ca^2+^ sensitive pool of filaments exists, our other results imply that it does not influence dynein accumulation and centrosome reorientation. It is also possible that the discrepancies in question reflect species-specific effects; we used primary murine T cells while the previous study used human material [20]. Recently, it was shown that human, but not mouse, NK cells require Ca^2+^ signaling for lytic granule exocytosis [30]. Hence, it is not implausible that synaptic F-actin dynamics might be governed by different signaling constraints in human than in mouse T cells.

Previously, we showed that synaptic accumulation of DAG and nPKCs downstream of PLCγ is required for centrosome polarization [7,8]. Our present work extends these observations by demonstrating that this same pathway also controls F-actin clearance and dynein accumulation. The nPKC substrates responsible for mediating these effects remain to be identified. Myristoylated alanine rich PKC substrate (Marcks) and the related Marcksl1 are intriguing candidates in this regard. Marcks family proteins are thought to couple the plasma membrane to the F-actin cytoskeleton [31]. PKC-mediated phosphorylation releases Marcks and Marcksl1 from the membrane and could therefore alter cortical F-actin architecture. We have found that TCR stimulation induces Marcksl1 depletion from the IS [8], and it will be interesting to investigate whether loss of one or both proteins disrupts F-actin clearance, dynein recruitment, and centrosome reorientation. PKC independent DAG signaling is also likely to play a role in these events. Indeed, the fact that unpolarized DAG signaling (via PBDU) disrupted synaptic F-actin architecture more strongly than did PKC inhibition (via Gö6983) in our hands implies that PKCs are necessary but not sufficient for T cell polarization. Furthermore, recent studies have implicated PIP_2_ depletion in synaptic F-actin clearance, highlighting the importance of PLCγ not only as a generator of DAG but also as a consumer of PIP_2_ in this context [9,32]. Deciphering the choreography of these diverse PLCγ dependent pathways will be an important step toward understanding the complex biology of the IS and of immune cell-cell interactions more generally.

## Materials and Methods

### Ethics Statement

The animal protocols used for this study were approved by the Institutional Animal Care and Use Committee of Memorial Sloan-Kettering Cancer Center.

### T Cell Culture and Retroviral Transduction

CD4^+^ T cells derived from transgenic mice expressing the 5C.C7 T-cell receptor were stimulated by coculture with B10A splenocytes at a ratio of 1:10 in RPMI medium containing 10% FBS and 5µM moth cytochrome c_88-103_ (MCC) peptide. 30 U/mL of IL-2 was added to cells 16 h following T cell activation. Subsequently, cells were split as needed into RPMI containing 30 U/mL of IL-2. T cells were retrovirally transduced with fluorescent probes as previously described 72 hours after the initiation of the T cell culture [7]. 24 hours post-transduction, cells were placed under puromycin selection (5 µg/mL). After an additional 48 h, transduced T cells were isolated by centrifugation over histopaque (Sigma).

### Signaling Probes

Lifeact constructs have been described [5,7]. The full-length coding sequence of centrin 2 was amplified from RNA derived from 5C.C7 T cell blasts and ligated into a murine stem cell virus (MSCV) retroviral expression vector upstream of and in-frame with Tag-RFP-T. Cytoplasmic dynein subunits were cloned from RNA isolated from 5C.C7 T cell blasts. All light intermediate chains (Dync1li1, Dync1li2 and Dync2li1), Tctex-1 light chain (Dynlt1), roadblock family light chains (Dynlrb1 and Dylrb2) and LC8 family light chains (Dynll1 and Dynll2) were inserted into pMSCV upstream and in-frame of either GFP or Tag-RFP-T. Tctex-3 light chain is inserted into MSCV retroviral vector downstream and in-frame of GFP. For visualization of the Dynactin complex and Lis1, we amplified the p50 subunit (dynamitin) of Dynactin and Lis1, respectively, from RNA isolated from 5C.C7 T cell blasts, and ligated them into pMSCV upstream and downstream of GFP, respectively.

### Photoactivation and TIRF Imaging

T cells expressing probes of interest were transferred into minimal imaging medium lacking phenol red and then attached to glass coverslips coated with NPE-MCC-I-E^k^ (125 ng/ml), nonstimulatory I-E^k^ containing a peptide derived from hemoglobin (amino acids 64-76; 3 µg/ml) and an antibody against H-2K^k^ (to encourage T cell attachment to the surface; 0.5µg/ml, BD Biosciences) [7]. Time-lapse images were recorded every 5 s for a total of 7 minutes using an inverted fluorescence microscope (Olympus) fitted with a 150× objective (1.45 NA). Fluorescent probes were all visualized using TIRF illumination except the centrosomal probes (tubulin and centrin), which were imaged in epifluorescence mode. GFP and RFP excitation was achieved using 488 nm and 561 nm lasers respectively. T cells were photoactivated with a 1.5 s UV pulse after the 14^th^ time point. UV irradiation of defined regions was performed using a Mosaic digital diaphragm system (Photonic Instruments) attached to a mercury lamp (Olympus). Small-molecule inhibitors and pharmacological agents were added to the medium above the cells as 200x stocks in DMSO; final concentrations were 50 nM Gö6983 (BioVision), 1 µM PDBU (Cell Signaling), 100 nM and 1 µM jasplakinolide (EMD Millipore), 30 µM nocodazole (MedChem Express), 1 µM taxol (MedChem Express), 50 µM ciliobrevin D (EMD Millipore), and 1 mM EGTA with 4 mM MgCl_2_ for calcium blockade.

### Lipid Bilayers and TIRF Imaging

Supported lipid bilayers containing streptavidin and biotinylated proteins were prepared as previously described [10]. To activate 5C.C7 T cells, lipid bilayers were coated with 1 µg/ml of biotinylated MCC-I-E^k^ and 1 µg/ml ICAM-1. T cells were incubated on stimulatory bilayers for 15 min at 37°C, followed by fixation in 2% paraformaldehyde for 10-15 minutes at room temperature (RT). For phalloidin staining, fixed cells were permeabilized using 0.5% Triton X-100 for 5 min, followed by incubation with PBS solution. Cells were then incubated with 1 µg/mL Alexa Fluor 594-labeled phalloidin in PBS for 1-2 hours at RT. T cells were imaged by TIRF microscopy with a 60× objective lens using a 561 nm laser. Small molecule inhibitors and pharmacological reagents were mixed with the T cells prior to their addition to the lipid bilayers; final concentrations were 50 nM Gö6983 (BioVision), 1 µM PDBU (Cell Signaling) and 1 mM EGTA with 4 mM MgCl_2_ for calcium blockade.

### Image Analysis

For photoactivation experiments, images were analyzed using SlideBook (Intelligent Imaging Innovations) and Microsoft Excel and graphed using Prism (GraphPad). Centrosome polarization was quantified by measuring the distance between the centrosome and the center of the UV irradiated region at each time point. Depletion or accumulation of fluorescent signaling probes at the irradiated region was quantified by calculating the mean fluorescence intensity (MFI) within the region for each time point. The MFI was normalized using the first ten time points prior to photoactivation, following background correction. T cell spreading on lipid bilayers was determined by calculating the area of the Alexa Fluor 594-labeled phalloidin signal. To calculate the ring ratio, the background-corrected mean fluorescence intensity (MFI) was compared to the MFI at center of the IS as previously described [10]. Ring ratios calculated from two perpendicular line scans were averaged to yield a single value for each cell.

### Immunoblot analysis

For T cell activation, 5C.C7 T cells were preincubated with 100 nM or 1 µM jasplakinolide or DMSO for 15 minutes at 37°C. Following treatment, T cells were mixed with polystyrene beads coated with 1 µg/mL of MCC-I-E^k^ and 1 µg/mL ICAM-1 and incubated at 37°C. At various time points, cells were collected and lysed in cold lysis buffer containing 50 mM TrisHCl, 0.15 M NaCl, 1 mM EDTA, 1% NP-40, 0.25% sodium deoxycholate, phosphatase inhibitors (1 mM NaF and 0.1 mM Na_3_VO_4_), and protease inhibitors (cOmplete mini cocktail, EDTA-free, Roche). Activation of PI3K and MAP kinase signaling was assessed by immunoblot for pAkt (Phospho-Akt (Ser473) Ab; Cell Signaling Technology) and pErk1/2 (Phospho-Thr202/ Tyr204; clone D13.14.4E; Cell Signaling Technology), respectively.

## Acknowledgements

We thank C. Firl and A. Kepecs for technical support and members of the Huse lab for assistance and advice. This work was supported in part by the NIH (R01-AI087644 to M. H., P30-CA008748 to MSKCC).

## Author Contributions

Conceived and designed the experiments: ES XL MH. Performed the experiments: ES XL. Analyzed the data: ES XL. Wrote the paper: ES MH.

